# Structural correlates of human muscle nicotinic acetylcholine receptor subunit assembly mediated by δ(+) interface residues

**DOI:** 10.1101/2020.06.11.145466

**Authors:** Max Epstein, Susan Maxwell, Thomas J. Piggot, David Beeson, Isabel Bermudez, Philip C. Biggin

## Abstract

Muscle nicotinic acetylcholine receptors are a class of heteropentameric ligand-gated cation channels with constituent subunits adopting a fixed stoichiometric arrangement. The specific amino acid residues that govern subunit ordering are however, only partially understood. By integrating all-atom molecular dynamics simulations, bioinformatics, two-electrode voltage clamp electrophysiology and ^125^I-α-bungarotoxin assays of chimeric nAChR subunits, we identify residues across the extracellular, transmembrane and extended M4 helix of the δ subunit that make structural signatures that contribute to intransigent assembly rules. Furthermore, functional differences observed in α_2_δ_2_β receptors can be rationalized by changes in dynamical behavior that manifest themselves at the agonist binding site.

## Introduction

Nicotinic acetylcholine receptors (nAChRs) are a class of pentameric ligand-gated ion channel (pLGIC) that come in a number of different combinations given their particular subunit composition. This composition effects overall biophysical characteristics and pharmacology, with different stoichiometries appearing preferentially in different tissue types [1]. NAChRs are comprised of one to four different subunits selected from an overall repertoire of 17 possible subunits in vertebrates [1]. End-plate nAChRs consist of α, β, δ and ε subunits in the case of adult muscle and α, β, δ and γ for foetal or denervated muscle, arranged in a specific order (**Fig. 1a**), that was first established by the work on Unwin and more recently confirmed by Hibbs and colleagues [2]. In the wild-type adult muscle nAChR, the orthosteric binding sites are located at α/δ and α/ε interfaces. The α subunit is typically referred to as the principal subunit (indicated by “+”), whilst the partner subunit is referred to as the complementary subunit (“-”).

**Fig. 1.**
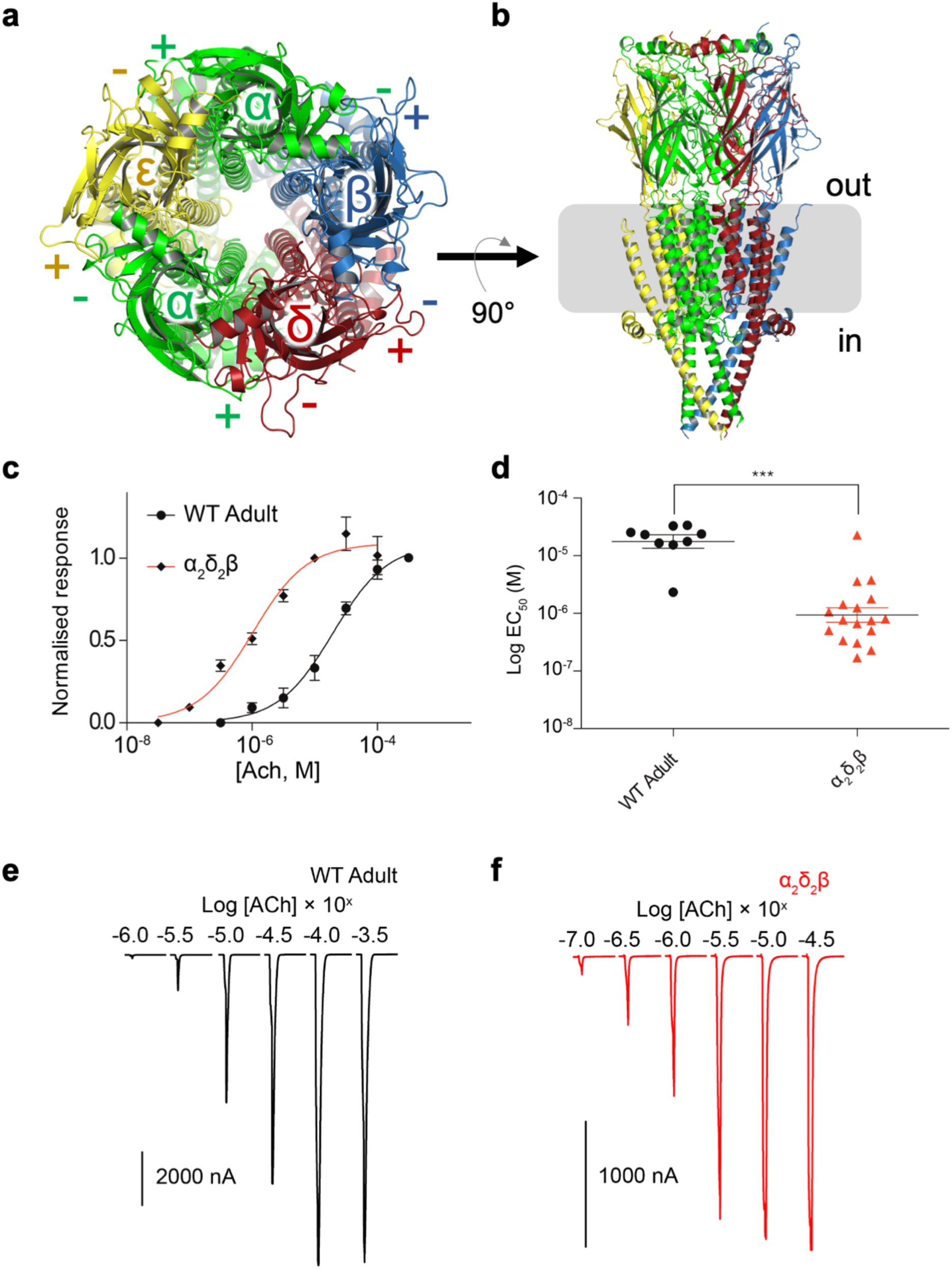
Electrophysiological measurements on WT and α_2_δ_2_β receptors. **a** Homology model showing arrangement of α, β, δ, and ε subunits coloured green, blue, red and yellow respectively, in the human muscle adult nAChR viewed from the synapse. **b** Model rotated 90° with the approximate location of the membrane indicated by a grey rectangle. **c** Normalized log concentration (in M) response curve of ACh at WT and α_2_δ_2_β stoichiometry nAChRs. **d** Spread of EC_50_ values from ACh concentration response curves with mean and 95% confidence intervals shown and p < 0.0001 denoted by asterisks. **e** Representative current traces of WT (black) and **f** α_2_δ_2_β receptor (red).

The invariant stoichiometry and large overall subunit repertoire implies a high degree of allostery that can be attributed to the requirement of individual subunits to be at defined locations with respect to the overall quaternary structure. Indeed, the nAChR has been shown to act as a so-called ‘allosteric machine’ [3]. The effect of any given subunit on muscle nAChR function can therefore be thought of as context dependent [4] and that if complementary orthosteric subunit faces (i.e δ and ε) were exchanged, whilst preserving all other structural aspects of the receptor, the overall receptor biophysics would differ from wild type. To what extent this is the case however is not understood and in order to answer this question, the full set of molecular determinants for specific stoichiometric assembly needs to be elucidated first.

Studies in GlyRs [5,6], GABA_A_Rs [7], 5HT_3_Rs [8] and nAChRs [9-16] provide clues as to the individual amino acid residues responsible for governing receptor assembly, expression efficiency, and the underlying mechanisms for oligomerization. However, not all of the molecular determinants that control subunit order in the human nicotinic acetylcholine receptor have yet been found. In previous work [15], mouse β/*γ* chimeras were used to determine the location of β(-) face residues important for assembly. Additionally, the molecular determinants for the mouse *γ* subunit at α(-) and α(+) interfaces were shown to reside at the extracellular domain (ECD). What *δ*(+) residues are responsible for forming the apparently exclusive interface with the β(-) face have yet to be specifically determined.

Previous studies of non-human muscle nAChR subunits in heterologous expression systems [17-19] and *in vivo* [20] have shown a certain degree of flexibility for the arrangement of muscle subunits that is limited predominantly to the so-called ‘double-δ’ stoichiometry but with evidence of a larger number of arrangements in mice [21] and rat [22] as well as inter-species variation with regards to expression efficiency [23]. There is currently a lack of evidence for variant stoichiometries for human subtypes.

By examining the propensity for variant muscle nAChR stoichiometries of heterologously expressed human cDNA subunits, we have elucidated permitted interfaces. The inability to express α_2_ε_2_β and α_2_γ_2_β stoichiometries in contrast to α_2_δ_2_β alluded to the presence of molecular determinants responsible for WT arrangements arising from the δ(+)β(-) interface.

Supplementation of this initial electrophysiological assessment of permitted interfaces with extensive all-atom molecular dynamics simulations and bioinformatic analysis by way of a multiple-sequence alignment allowed a more detailed description of the specific molecular determinants responsible for this invariant interface. This was then validated by ^125^I-α-BuTx binding to chimeric subunits transiently transfected in HEK293 cells.

Furthermore, we sought to characterise the biophysical properties of the human αδβ nAChR that we obtained under experimental conditions and determine a structural explanation for its differing behaviour to WT by comparing simulations of apo and acetylcholine (ACh) bound states of human adult muscle nAChR. The simulations suggested little change in the ACh binding modes as expected. However, we also observed a distinct increased mobility of loop-C at the αδ interface in the apo state, compared to the αε interface. Taking the large (18.75 fold) reduction in EC_50_ for the α_2_δ_2_β receptor compared to WT with the observations from simulation, we hypothesise that increased C-loop motility at the agonist binding site could allow for a less occluded binding site and would facilitate the recognition of ACh during the initial diffusive stage of agonist binding.

## Methods

### Homology modelling

Full length sequences of adult human end-plate nAChR subunits were aligned with the target sequences of α_3_ and β_4_ nAChR subunits which correspond to the subunits present in the template model (6PV7) [24]. Additional phylogenetically related subunits were included to improve confidence in the multiple sequence alignment, which was then performed using MUSCLE [25] before manual editing to obtain the final alignments (**SI Fig. 1**). In the first instance a single model was generated. This model was further refined by generating 10 additional models for three different non-conserved loop regions sequentially. Modeller [26] was used to build the models. Specifically, this included residues 562-572, 969-977 and 1768-1776 situated on δ, β and ε subunits respectively and numbered according to sequential .pdb residue numbering. At each step of the loop refinement procedure, models were carried forward for additional loop refinement based on the best molpdf score. In total, 31 models were generated before arriving at the final template structure. The intracellular domain was not included as it was not resolved in the template structure.

### Docking

ACh was docked using a grid box spacing of 1.0 Å with the docking box centered on the aromatic box for both orthosteric interfaces. Exhaustiveness was set to 8 and docking calculations performed in Vina 1.1.2 [27]. For all ligands, Gasteiger charges [28] where added and non-polar H atoms merged using Autodock 1.5.6 [29]. Docked poses were selected based on the proximity of the choline moiety to the centre of the aromatic box.

### MD simulations

The homology model was embedded in a 100% POPC bilayer using inflategro.pl [30] before being solvated using a TIP3P water model [31]. Na^+^ and Cl^-^ ions were added to neutralise the system before additional ions to make the system up to a final concentration of 0.15 M. This system was then energy minimised via the steepest descent algorithm (step size 0.01 nm, max number of steps = 50000). Electrostatics where calculated using the Particle mesh Ewald algorithm [32] with electrostatic cut off set to 10 Å, a cubic interpolation and fourier spacing of 10 Å. A 10 Å cutoff for the Van der Waals interactions was implemented. H-bonds were constrained using LINCS [33] to allow for a 2 fs time-step. Amber ff99SB-ILDN was used as the protein forcefield [34] and lipids were modelled using Slipids [35]. Each system was equilibrated in the NPT ensemble for 1 ns using semi-isotropic Berendsen pressure coupling [36] (tau = 1.0) and temperature coupling with V-rescale [37] (tau = 0.1) at 310 K. Backbone atoms were restrained with 1000 kj/mol/A^2^ force constant in order to allow for solvent and lipid molecules to equilibrate around the protein. Due to significant expansions in the z-axis after this short equilibration step, all water molecules were removed and periodic boundary conditions reduced along this axis before re-solvating and re-ionising the resulting system to decrease excessive buffer and improve overall sampling. The resulting system was then equilibrated for a further 5 ns in the same thermodynamic ensemble with holonomic constraints applied to h-bonds using the LINCS algorithm and position restraints removed to allow proper equilibration of the protein. Production runs were carried out with the Nosé-Hoover [38] (tau t=0.5) and Parrinello-Rahman (tau p=1.0) thermostat and barostat for 100 ns each. All simulations were performed using gromacs [39].

### Multiple Sequence Alignment and Contact Matrix analysis

For the identification of potential residues involved in arrangement-an MSA of mammalian δ, ε and *γ* subunits was generated (**SI Fig. 2**). Candidate positions were selected based on whether at a specific point in the alignment, both *γ* and ε residues were conserved whilst δ residues differed. A pair-wise protein-protein Cβ atom contact prediction with a cut-off distance of 8 Å was selected according to the CASP criteria [40] and generated using MDanalysis [41]. Specific residues are numbered according to order of appearance in respective uniprot fasta files whilst disregarding signal peptide sequence. All residue numbers are labeled according to the mature peptide sequence herein.

**Fig. 2.**
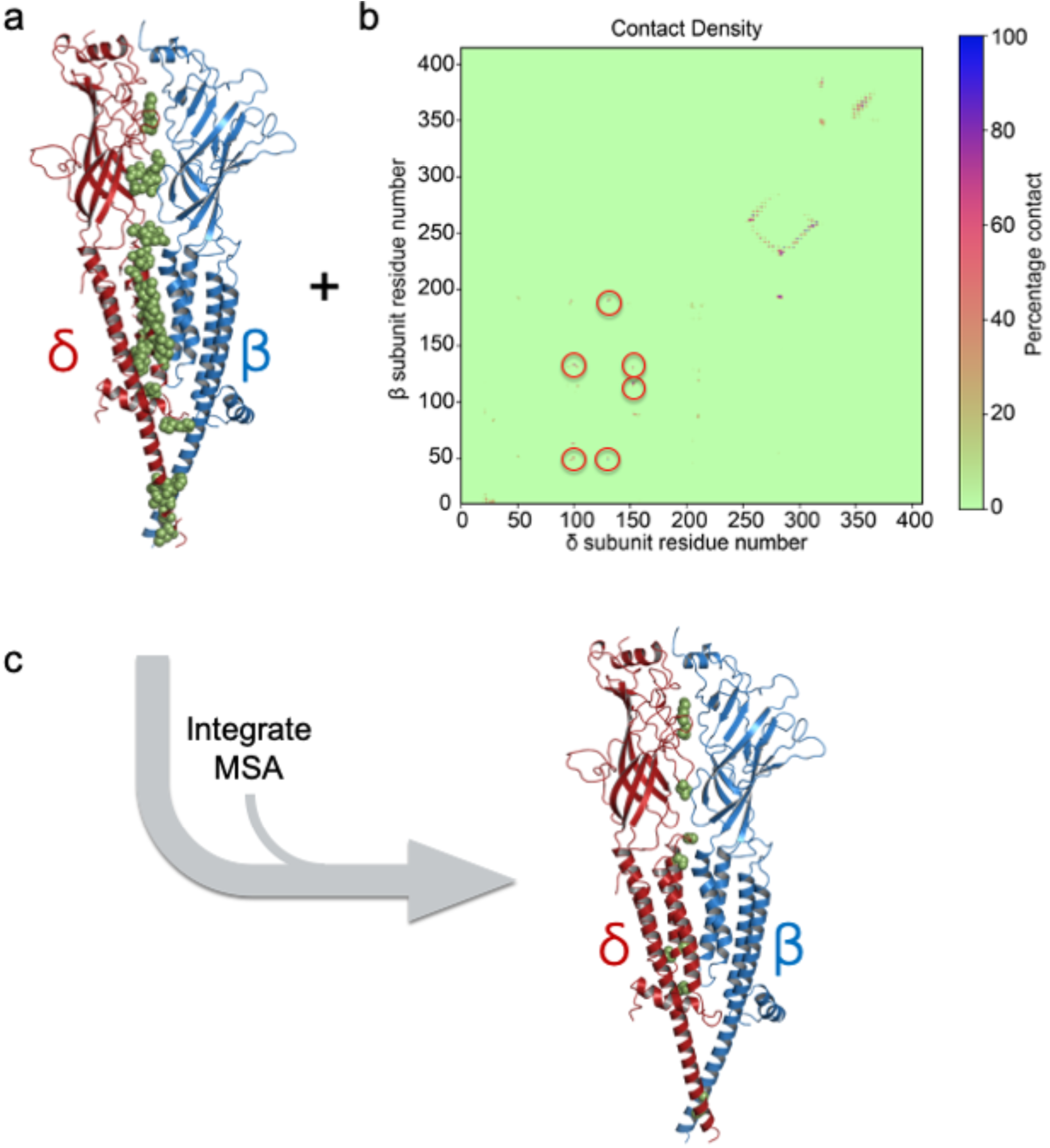
Identification of important contacts. **a** Representative snapshot from apo simulations. δ and β subunits shown in red and blue respectively. Green sphere representation denotes all δ subunit residues who’s Cβ atoms or Cα for glycine came within 8 Å of the β subunit over 70% of aggregate simulation time. α and ε subunits have been removed for clarity. **b** Pairwise residue contact matrix with red circles used to clarify single contact points in the ECD. **c** Remaining residues (green spheres) after filtering the contact matrix by MSA criteria (see main text for definition).

### Molecular Biology

Wild type human muscle cDNA subunits {Beeson, 1993 #9554} were subcloned into pcDNA3.1 (Invitrogen). The δ-ε-ε chimera was generated by PCR with overlap extension [42] (**SI Table 1**). Additional chimeric constructs were synthesised by GeneArt (Regensburg, Germany). Full-length sequences of mutant subunit cDNAs were verified by sequencing (Eurofins-MWG, Germany). Specifically, δ/ε domain swaps occurred betweenδ residue numbers 231 for the ECD-TMD boundary, 320 for M3-MX & ICD boundary and 418 for the ICD-M4 boundary with the corresponding aligned residues according to **SI Fig. 1**.

**Table 1.** ANOVA with Tukey’s multiple comparison test for the surface expression of different δ subunit chimeras.

| Construct | Significance |
| --- | --- |
| WT vs $\alpha\delta\beta$ | *** |
| WT vs $\delta\epsilon\epsilon$ | *** |
| WT vs $\delta\delta\epsilon$ | *** |
| WT vs $\delta\epsilon\delta$ | *** |
| WT vs $\epsilon\delta\epsilon$ | *** |
| $\alpha\delta\beta$ vs $\delta\epsilon\epsilon$ | *** |
| $\alpha\delta\beta$ vs $\delta\delta\epsilon$ | *** |
| $\alpha\delta\beta$ vs $\delta\epsilon\delta$ | *** |
| $\alpha\delta\beta$ vs $\epsilon\delta\epsilon$ | *** |
| $\delta\epsilon\epsilon$ vs $\delta\delta\epsilon$ | *** |
| $\delta\epsilon\epsilon$ vs $\delta\epsilon\delta$ | ns |
| $\delta\epsilon\epsilon$ vs $\epsilon\delta\epsilon$ | ns |
| $\delta\delta\epsilon$ vs $\delta\epsilon\delta$ | * |
| $\delta\delta\epsilon$ vs $\epsilon\delta\epsilon$ | * |
| $\delta\epsilon\delta$ vs $\epsilon\delta\epsilon$ | ns |

\*\*\* < 0.0001, \* < 0.05, ns = not significant

### ^125^I-α-BuTx binding assay, cell culture and transfections

^125^I-α-BuTx binding assay, cell culture and transfections were carried out according to the protocol outlined in previous work [43] in which 2:2:1 ratios of α, chimeric and β subunits and α, δεγ and β subunits were transfected into HEK293 cells for chimeric and double-’δεγ’ stoichiometries. 2:1:1:1 ratios of α, δ, β, ε and α, δ, β, γ were transfected into HEK293 cells for WT stoichiometries. ^125^I-*α*-BuTx binding assays were done 2 days post-transfection with each assay performed in triplicate wells to ensure reproducibility. Results were normalized according to WT counts performed on the same day and seeded from the same original batch of HEK293 cells.

### Expression of nAChRs in Xenopus Oocytes

*Xenopus laevis* oocytes were purchased from Xenopus One (Michigan USA). Toad ovary dissection was done according to regulations set out by the UK Home Office. Xenopus oocytes were prepared as described by previous methods [44]. Wild type human nAChR α_1_β_1_δε subunit cDNAs were co-injected into the nuclei of oocytes at a 1:1:1:1 ratio and volume of 50.6 nl/oocyte, using a Nanoject Automatic Oocyte Injector (Drummond, Broomall, PA, USA). 2α_1_2δβ_1_, 2α_1_2εβ_1_, 2α_1_2*γ*β_1_ stoichiometries were injected at a radio of 2:2:1. 2 ng of cDNA was injected per oocyte. Post injection, oocytes were incubated for 2-5 days at 18 °C in modified Barth’s solution containing1 mM KCl, 2.4 mM NaHCO3, 88 mM NaCl, 0.3 mM Ca(NO3)2, 0.82 mM MgSO4, 0.41 mM CaCl2, 15 mM HEPES and 5 mg/l neomycin (pH 7.6).

### Electrophysiology and concentration response curves

All recordings were performed within 2 to 5 days after injection. Oocytes were placed in a recording chamber of 0.1 ml and perfused at 15 ml/min with Ringer’s solution of composition: KCl 2.8 mM, NaCl 150 mM, CaCl_2_ 1.8 mM, HEPES 10 mM; pH 7.4 adjusted with NaOH and NaCl to obtain the desired pH. The two-electrode voltage-clamp (TEVC) configuration was used to obtain current responses, using a holding potential of −60 mV. The automated HiClamp setup (Multi Channel Systems) was used to obtain electrophysiological recordings. 3 M KCl was used as the electrode solution with a small aliquot of mineral oil at the entrance of each electrode to prevent evaporation and the formation of salt crystals. All experiments were performed with electrodes that had resistances of less than 1 MΩ. Serial dilutions of ACh in Ringer’s solution was done on the day of use from frozen stock of 10 mM. Experiments were performed at room temperature. For all experiments, a 6 to 8 point log concentration-response curve (CRC) was obtained. Currents were normalised relative to ACh EC_100_ (1mM). Washes between ACh applications were 3 minutes long. Stabilisation of oocytes with ACh EC_100_ was done at the start of each experiment and intermittently after every two ACh applications in order eliminate run-up or run-down of responses from being included in subsequent data analyses.

### Data analysis

All CRCs were fit with to a non-linear regression, mono-phasic four-parameter Hill equation using Prism 5 software (Graph Pad, CA, USA). Individual EC_50_ values for electrophysiology experiments were plotted and analyzed by way of unpaired two-tailed t-test. A p value of less than 0.05 was deemed to be significantly different. ^125^I-α-BuTx binding assays were analysed by one-way ANOVA with Tukey’s post hoc test to control for multiple comparisons.

## Results

### The expression of α_2_δ_2_β nAChRs and lack of α_2_ε_2_β or α_2_γ_2_β nAChRs implies exclusive contacts at the δ(+)β(-) interface

2:2:1 ratios of αδβ, αεβ or αγβ nAChR subunit cDNAs were injected into Xenepous oocytes to assess whether these variable stoichiometries could express competent receptors. As no currents could be detected for αεβ and αγβ injections,1:2:1 ratios were also attempted - again with no detection of currents. α_2_δ_2_β receptors exhibited a statistically significant difference to WT EC_50_, with a resulting decrease in EC_50_ from 17.58 μM (upper 95% CI = 32.66 uM, lower 95% CI = 9.46 μM) (n=9) to 0.938μM (upper 95% CI= 1.73 μM, lower 95% CI = 0.51 μM) (n=17) (**Fig. 1**). In order to rule out the possibility of cell surface expression of non-functional receptors, ^125^I-α-BuTx binding assays were performed on these subunit ratios with no detectible expression on the cell surface. As α_2_γ_2_β and α_2_ε_2_β stoichiometries could not form functional pentamers, this indicated that either the δ subunit was somehow adapted to form an exclusive interface with the β(-) face and or that the β(-) interface prevents formation with γ and ε subunits.

### MD simulations and MSA integration point to specific residues on ECD, TMD and M4 extended helix of the δ subunit as important in forming exclusive contacts with β(-)

To obtain a more precise estimate of which residues in the δ(+) subunit contribute to the uniqueness of the δ(+)β(-) interface, we ran 10 x 100 ns all-atom molecular dynamics simulations of the adult human muscle nAChR. A pairwise contact matrix was employed to analyse each individual residues Cβ carbon atom distances between δ and β subunits (**Fig. 2**). A cutoff distance of 8 Å was selected according to the CASP criteria for protein-protein interaction distances. Residues that fell within this criteria for over 70% of aggregate simulation time (**Fig. 2b**) were then projected on to a snapshot from one of these simulations as shown in **Fig. 2a**. When 60% of aggregate simulation time is used to determine contacts, there was negligible change, with only δ H335 interactions no longer being detected.

In order to filter out spurious residues that reside at the δ(+)β(-) interface but are not important in determining the interface, an MSA of all uniprot approved sequences of mammalian δ, ε and γ subunit sequences was performed (**SI Fig. 2**). As ε and γ subunits occupy the same position in the global receptor WT topology it can be inferred that the residues that permit their particular positions are conserved with respect to one another and that corresponding δ subunit residues should also be conserved but not when compared to ε and γ. Residues that fit this evolutionary criteria and that were also detected by the contact matrix distance analysis were identified and mapped to the structure (**Fig. 2c**). Whilst intracellular domain residues were detected in the MSA that fulfilled this evolutionary criteria, the lack of structural template of which to model this domain on meant that they could not be detected in the contact analysis. Percentage contact time for each specific residue identified from the contact analysis and MSA integration was calculated (**SI Table 2**) to help manually check the data.

### ^125^I-α-BuTx assays on chimeric subunits confirm that δ(+)β(-) specific interface determinants occur across domains

In order to experimentally confirm that residues important for forming the δ(+)β(-) interface occur across domains, a series of chimeric δ/ε subunit constructs were made (δ_chimera_) (**Fig. 3a**) and injected in 2:2:1 ratios to investigate the role of three domains of the δ subunit were necessary for functional expression. The resulting chimeric receptors in transfected HEK293 cells and their cell surface expression were measured by ^125^I-α-BuTx binding to surface expressed chimeras (**Fig. 3**). A receptor comprised of purely WT δ subunits (ie a receptor with the following composition: α_2_δ_2_β) shows a reduced surface expression (**Fig. 3b**) with respect to WT (ie α_2_δεβ), but surface expression is still robust and can be explained by the loss of favourable contacts between δ(+) and α(-). It should be noted that as both *δ* and ε subunits are tolerated between α subunits, the use of double chimeras will not preclude the analysis of δ(+)β(-) interface. However, it may result in reduced overall surface expression. The δ subunit chimera constructs δεε, δεδ and εδε show little to no surface expression (**Fig. 3b**) and are not statistically significantly different from one another (**Table 1**). Receptors with the δδε chimera however, are capable of reaching the surface of the cell (**Fig. 3b**) and are statistically significantly different from δεε, δεδ and εδε containing constructs (**Table 1**). The δδε construct containing receptors are also statistically significantly different from the αδβ nAChR (α_2_δ_2_β) indicating that the intracellular domain is not critical in forming δ(+)β(-) interfaces, but does however promote increased cell surface expression efficiency. Overall, the expression of receptor with the δδε construct and the lack thereof for the receptors containing the δεδ and εδε and δεε constructs, indicates that both extracellular and transmembrane domains contain the molecular signatures for enabling δ(+)β(-) interface formation.

**Fig. 3.**
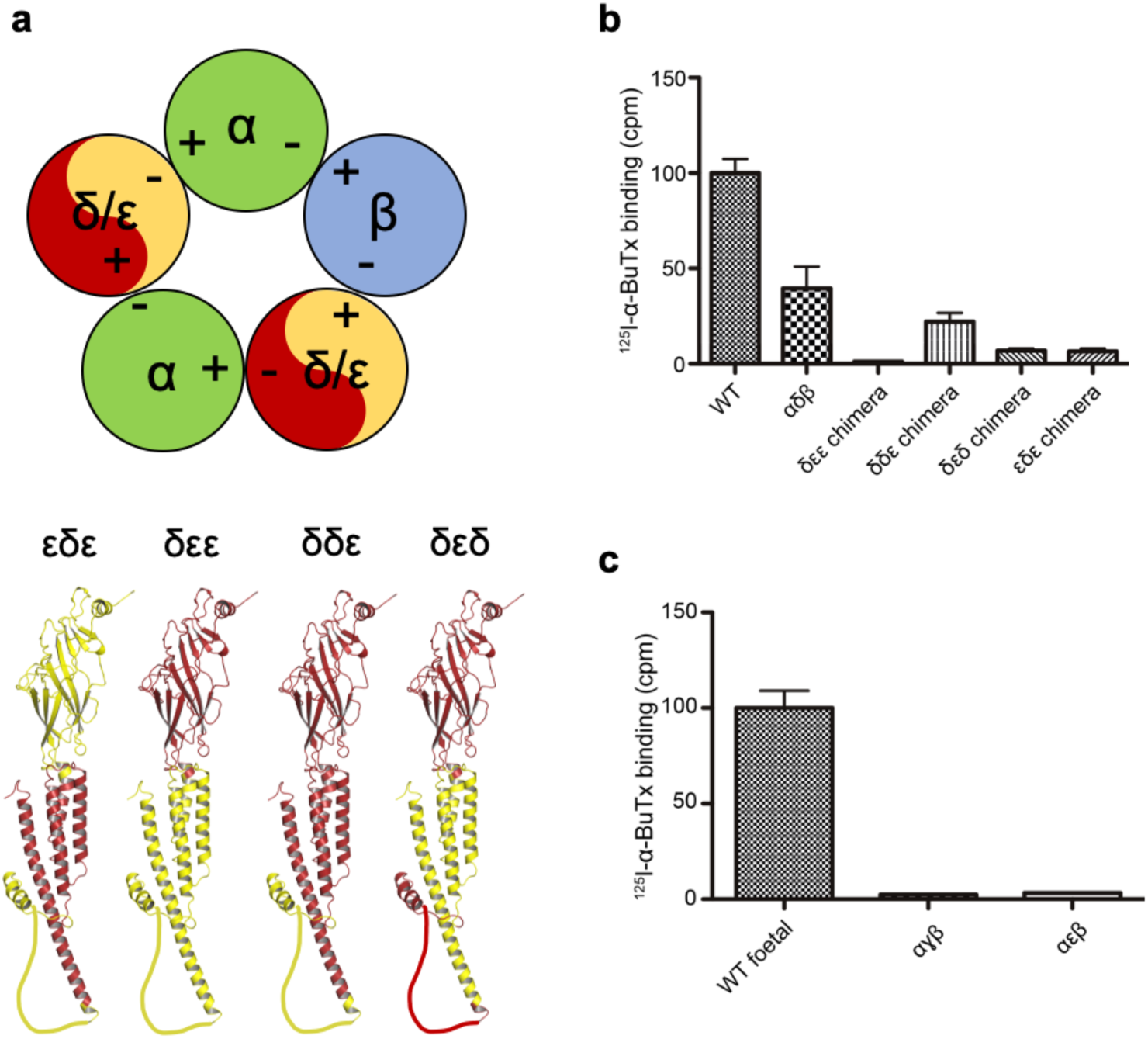
Chimeric construct strategy. **a** Schematic of constructs used for ^125^I-α-BuTx assay and associated models of each chimeric subunit, with ε segments in yellow and *δ* in red. **b** ^125^I-α-BuTx assay means (normalised to WT counts) with associated 95% confidence interval. WT n=12, αδβ n=6, δεε n = 6, δδε n = 6, δεδ n=6, εδε n=5. **c** ^125^I-α-BuTx data for αγβ (n=3) and αεβ (n=3) normalised to WT foetal receptor (n=2).

### δ**K173, δN119 and δS297 participate in interface contacts via hydrogen bonding with βS134, βQ62 and βN251 respectively**

MD analysis of specific residues highlighted from the integrated contact matrix analysis showed that δK173 is capable of participating in hydrogen bonding interactions with the proximal β face residue S134, with a peak observed at around 5 Å (**Fig. 4a**). There exists an additional peak at around 7 Å that would be indicative of non-interacting residues. Indeed, the overall span of this histogram indicates a high degree of flexibility for these residues. The ability of *δ*N119 was also further assessed to form hydrogen bonds with neighbouring β subunit residue Q62 as both residues Cβ atoms were within 8 Å of each other as determined from the contact matrix analysis. As both residues contain hydrogen bond donor and acceptor atoms, density histograms for each pair was plotted (**Fig. 4b**). Major residue-residue interactions are mediated via the δN119 carbonyl group and βQ62 amino group. While δN119 amino - βQ62 carbonyl interactions are possible, they represent a minority of the hydrogen bond interactions between this residue pair. A further interaction is found between at the top of the TMD between the sidechain OH atom of δS297 and the sidechain oxygen of βN251 (**Fig. 3c**).

**Fig. 4.**
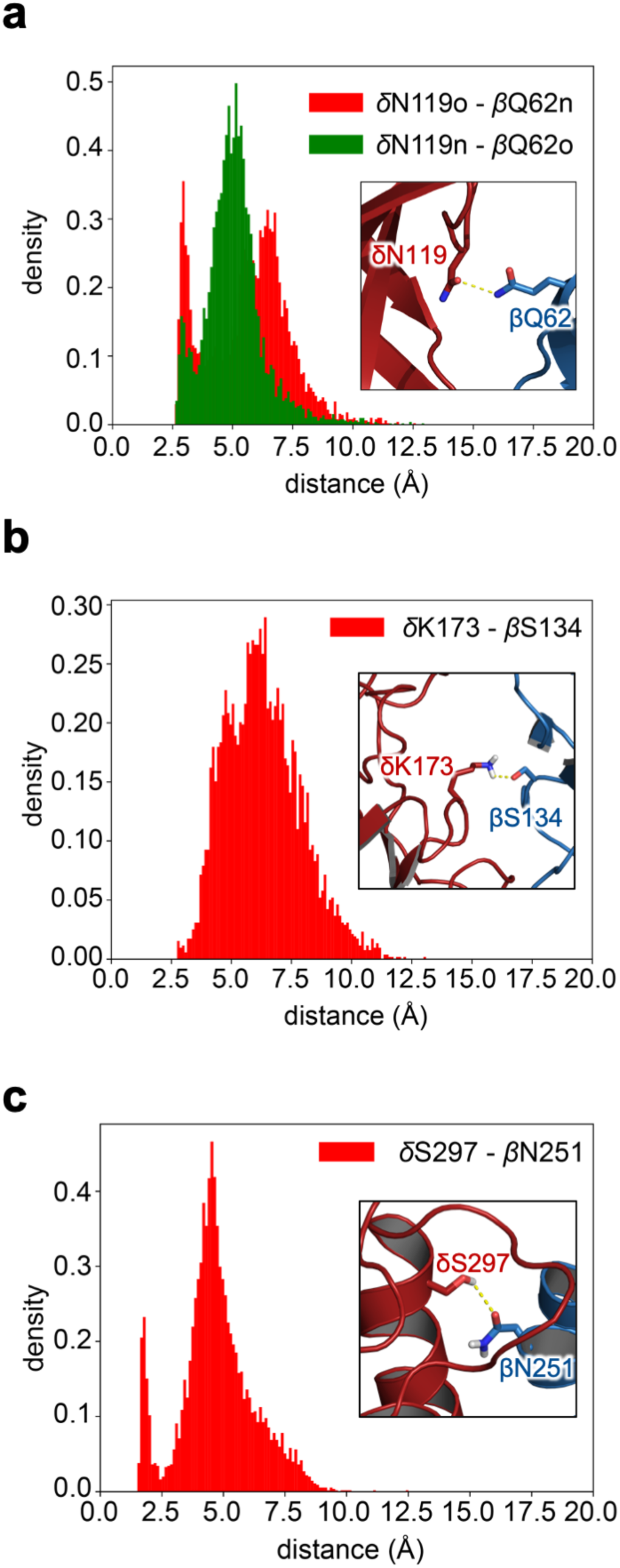
Density histograms of ECD δ(+)β(-) contacts. **a** Distance between each nitrogen and oxygen atom pair between δN119 and βQ62. **b** Distance between δK173 and βS134 nitrogen and oxygen atoms. **c** Distance between δS297 and βN251 nitrogen and oxygen atoms. Inserts are representative snapshots from simulations with δ in ‘fire-brick’ red cartoon representation and β in ‘sky-blue’ cartoon representation. Amino acids of interest are shown in the stick representation, with their oxygen and nitrogen atoms coloured red and blue respectively.

### C-loop dynamics are not equivalent at respective agonist-binding interfaces

Simulations were also used to further understand the shift in the EC_50_ that arises from a double δ receptor (**Fig. 1**). 10 additional 100 ns MD simulations were performed with ACh docked into both agonist binding sites and C-loop motions in apo and ACh-bound states compared. Comparison of the binding mode of ACh at both interfaces revealed no differences (**Fig. 5a**) in terms of the interaction of ACh with the binding pocket and is consistent with previous simulation reports of ACh dynamics in nAChRs [45]. The overall mode of interaction (**Fig. 5b**) is also preserved throughout all simulations. When ACh is bound, the C-loop mobility, as assessed by the variation of distance between the C-loop vicinal cysteines (Cys237 and Cys238) and the benzene ring of Trp194, does not differ appreciably between interfaces, although a negligible peak can be observed near 17.5 Å for the αδ interface. However, for the apo state, the mean position of the C-loop at the αδ interface is slightly shifted towards larger values, at around 11 Å, compared to approximately 10 Å for the αε interface. This is also coupled with a pronounced additional peak, observed at around 15 Å, suggesting that the C-loop can adopt a new, more open, state at the αδ interface.

**Fig. 5.**
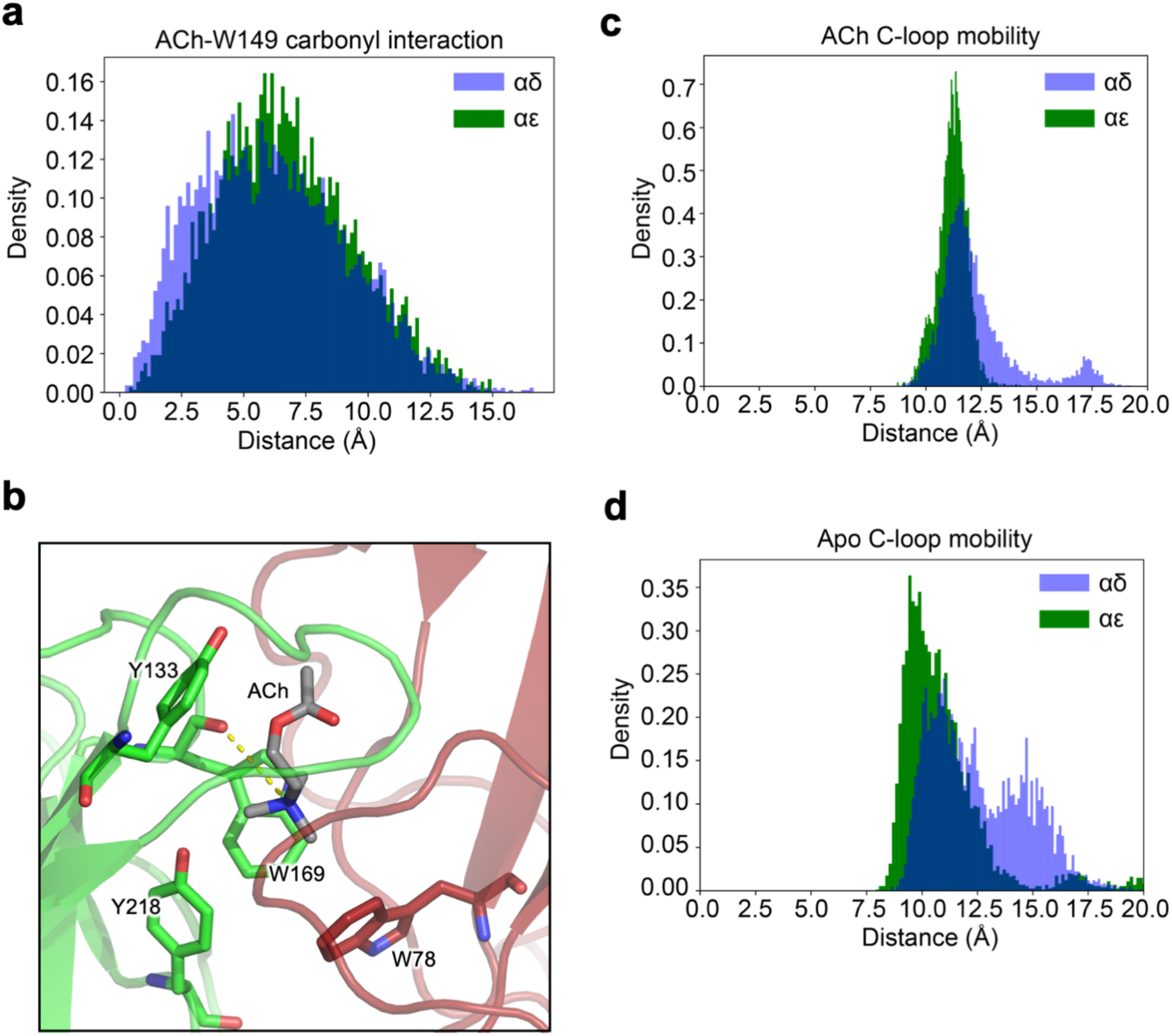
C-loop dynamics. **a** Density histogram of ACh ammonium nitrogen and W149 carbonyl oxygen at αδ and αε interfaces as exemplified by the snapshot in **b** with the distance between W149 carbonyl and ACh ammonium nitrogen displayed as a dashed yellow line. **c** Density histograms between centre of geometry of C-loop disulphide sulphur atoms and centre of geometry of benzene moiety of W149 α subunit at both ACh and (D) apo state nAChR.

## Discussion

The ability of nAChR subunits to assemble in variable stoichiometries expands the already large functional diversity of this receptor family. The muscle nAChR subtype is especially interesting in this regard, not only on account of the developmental switch that occurs during neuromuscular maturation [46,47], but also from the apparent intransigence of subunit ordering. It appears that ε/γ and δ subunits have evolved to occupy a fixed position in the global receptor topology despite the fact that if swapped for one another - both agonist binding sites would be preserved. This implies that the pseudo-symmetrical architecture of the receptor can allosterically effect each orthosteric site in unique ways and that the overall contributions to overall receptor biophysics arising from individual subunit interfaces depends on their context with regards to all other subunit positions. In order to obtain experimental evidence of this however, the molecular determinants at the δ subunit responsible for forming an interface with the β subunit must be first be elucidated. As all previous studies focus on non-human subtypes and given the large inter-species diversity commonly observed between specific nAChR stoichiometries [23] - and especially demonstrated by differences in response to ACh [48-51] this provided further justification for the study outlined herein. Additionally, while regions on both α inter-subunit interfaces that govern assembly have been identified for mouse muscle nAChRs [15]. How δ is adapted to exclusively form interfacial contacts with the β subunit in the context of a pentamer in human subtypes has not been shown. While attempts to delineate interfacial components have been addressed via cryo-EM structures of *Torpedo* ACh receptors [2,52], this does not precisely pinpoint assembly specific residues. Retrospective evaluation of the recently published *Torpedo* structure [53], 6UWZ, also shows species specific variation at assembly identified positions (**SI Fig. 3**).

Injection of 2:2:1 ratios of respective subunits for α_2_ε_2_β, α_2_δ_2_β and α_2_γ_2_β stoichiometries only yielded currents for α_2_δ_2_β. This pointed to residues on δ(+)β(-) face as the determinants for stoichiometric invariance. Furthermore, α_2_δ_2_β shows a large shift in EC_50_ for ACh in agreement with previous results for bovine muscle receptors [18]. Given this and previous data, a subunit order of αδαδβ (clockwise viewed from the synapse) is completely consistent. However, we cannot at this stage, completely rule out other alternatives. That would require the use of concatemers [54, Mazzaferro, 2014 #6892], a technique that has recently been used to show that loose subunits can lead to heterogenous receptor populations, although this was for α4β2α5 receptors [55]. Nevertheless, due to the robust currents recorded, no change in hill slope compared to WT and the requirement of a β subunit for cell surface expression [17,19], a reasonable and parsimonious assumption is that this receptor adopts a αδαδβ ordering.

Given that pLGIC ICDs can assemble independently of the rest of the receptor [8] and that functional channels can be obtained without intracellular domains [56], it is unsurprising that our δδε chimeric subunit can still form functional pentamers. The ICD is also known to participate in interactions with cytoskeletal proteins and in the case of the δ subunit, is predicted to be mostly unfolded [57]. This makes it very unlikely to participate in determining subunit ordering to any extent. The δδε chimeric construct also leads to decreased expression with respect to WT and α_2_δ_2_β, therefore, the ICD is important for expression efficiency but not for forming specific arrangements. Furthermore, mutation to δεε, causes complete ablation in expression. Comparing this with δδε indicates that δ transmembrane domain (TMD) residues are involved in some capacity for forming specific arrangements.

It would appear however that from the lack of expression for the εδε chimeric construct that residues identified on the δ ECD are also important in forming specific arrangements. This demonstrates that the molecular determinants for forming the exclusive δ(+)β(-) interface are not confined to a single domain and is particularly interesting given that previous determinants of assembly have been show to reside at ECD ‘assembly boxes’ [58,5].

From the MD simulations, hydrogen bond interactions between δN119 and βQ62 were observed, with well-defined peaks (**Fig. 4a**). Hydrogen bond donor and acceptor moieties for both residues do not seem to interact simultaneously which would theoretically increase the strength of this part of the interface, however the definition of peaks would be indicative of stable interactions overall for these residues. Hydrogen bonding was also observed between residues δK173- βS134 (Fig. 4B) with a broad overall histogram indicative of high residue flexibility. An identifiable peak at 5 Å probably corresponds to hydrogen bonding interactions, whilst the major peak occurred closer to 7 Å. The proximity of these residues to the solvent exposed region of the ECD funnel as well as the high level of flexibility would indicate sub-optimal interactions despite δK173 residing at the same position as conserved γ residues shown to be essential for forming γ(+)α(-) interactions in mice nAChRs [15]. It should also be noted that as well as the favourable interactions mediated by δ(+) interface residues responsible for allowing WT stoichiometry to occur, unfavourable interactions between homologous residues of the ε(+) face and β(-) face may also prevent ε(+)β(-) interfaces from occurring. δK173 and δN119 are replaced by threonine and isoleucine in the ε subunit and would represent the loss of potential hydrogen bond interactions (**SI Fig. 4**). To this end, homologous δ(+) and ε(+) residues could work in tandem with favourable and unfavourable interactions respectively to enable the fixed stoichiometry of WT human muscle nAChRs.

Residues δS297 and δA302 in the M2-M3 loop of the δ subunit show hydrogen bonding and hydrophobic interactions respectively with residues on the β subunit **(Fig. 4c and SI Fig 5a**). It is well established that the M2-M3 linker acts as a key structural motif in coupling agonist binding to gating [59,60]. Although mutations to ε and β subunits have been demonstrated to change the open time constant, no such change could be observed for homologous residues at the δ subunit [61]. Given this and from the data described herein, it is likely that part of this region could serve an alternative function and could be important in forming an optimal interface with the β(-) faces. While residues δE277 in the −1’ position and δT279 in the 1’ position of the M1-M2 loop were highlighted from the contact matrix analysis, the conservation of δE277 with β and α subunits, as well as its well-known role in determining ion selectivity [62] make it highly unlikely that it is involved in forming the δ(+)β(-) interface. Despite this, *δ*T279 is positioned within the inter-helical bundles of both subunits and is not conserved widely across muscle subunits (**SI Fig. 5b)**. The importance of inter-helical interactions in shaping overall receptor stoichiometry arising from intra-helical aromatic interactions has been previously eluded to [6]. H335 was also identified from the contact analysis, although with a significantly reduced overall percentage contact with the β-face compared to other residues (**SI Table 2**). Uncertainty around the protonation state of this residue also makes it difficult to determine its role, if any, in the δ (+)β(-) interface and results from the simulation for this residue should be taken with caution. Residues δP439 and δG442 were identified from the contact matrix analysis and reside toward the intracellular end of the extended M4 helix (**SI Fig. 5c)**. Given their physicochemical properties and solvent exposure, it is likely that they form highly favourable hydrophobic interactions with β subunit residues.

As a precursor to assessing the effect of δ and ε subunit position on receptor function and response to ACh, we wanted to first determine the extent of differences intrinsic to both α*δ* and αε interfaces. To do this, we performed additional MD simulations with ACh docked into both orthosteric sites of the WT adult nAChR and compared interfaces in the apo and ACh bound states. Whilst the binding mode of ACh was comparable between interfaces (**Fig. 5a**), loop-C dynamics was markedly changed when comparing between apo and ACh bound states (**Fig 5c, d**). The shifted density observed for the αδ interface in the apo state compared to the αε interface as well as an additional peak at around 15 Å indicates a non-occluded agonist binding site. This would translate mechanistically to a decreased *k*_*on*_ for ACh and could explain the shift in EC_50_ observed (**Fig. 1**). While this would manifest in differing K_d_ values for ACh at both binding sites, previous work in mouse muscle nAChRs has shown both interfaces to be functionally equivalent [45]. Despite this, there exist large variations in overall nAChR sensitivity to ACh across species [50,51,49]. In addition, the close proximity of the non-conserved δ loop-F to loop-C (**SI Fig. 6**) could feasibly cause changes in the response to ACh as it has been postulated that signal propagation post ACh binding occurs from the C-loop to the M2-M3 loop via the F-loop [63]. Differences in the contributions from both interfaces to gating have also been previously demonstrated [64] and given the 46.3% sequence identity between human δ and ε subunits, there are many structural opportunities for changes in ACh sensitivity.

## Conclusions

Using a combination of electrophysiology, bioinformatics and MD simulations, we have identified residues across the extracellular, transmembrane and extended M4 helix of the human nAChR δ subunit that contribute to the intransigent assembly of WT muscle nAChRs. This aligns with previous studies on the assembly of pLGICs and offers a new insight into the strict rules governing the specific arrangement of end-plate nAChRs. Additionally, we have shown that differences in EC_50_s between WT and α_2_δ_2_β nAChRs can be rationalised by the altered dynamics of the C-loop observed in MD simulations and which may influence ACh binding kinetics.

## Acknowledgements

We thank Dr Richard Webster and Dr Judith Cossins for useful discussions. M.E. is supported by an Engineering and Physical Sciences Research Council (EPSRC) Industrial Cooperative Awards in Science & Engineering (iCASE) studentship (voucher number 15220076) with the Defence Science and Technology Laboratory (DSTL). PCB thanks the BBSRC for funding (BB/S001247/1). This project made use of computation time on JADE (EP/P020275/1) via HECBioSim (http://www.hecbiosim.ac.uk), supported by EPSRC (grant no. EP/R029407/1). We thank Advanced Research Computing (Oxford) for additional computer time.

## Author contributions

P.C.B and M.E. conceived the study. M.E planned the simulation protocol, performed and analyzed the MD simulations. M.E. performed and analyzed the electrophysiology and I.B. supervised the electrophysiology. M.E. and S.M. performed the ^125^I-α-BuTx assays and M.E. analyzed the data. D.B. supervised the ^125^I-α-BuTx assays. P.C.B., T.J.P and M.E. interpreted the results. P.C.B and M.E. wrote the paper.

## Compliance with ethical standards

Dissection of *Xenopus laevis* was done in accordance with UK Home office regulations.

## References

1. Albuquerque EX, Pereira EFR, Alkondon M, Rogers SW (2009) Mammalian nicotinic acetylcholine receptors: from structure to function. Physiol Rev 89 (1):73–120. doi: 10.1152/physrev.00015.2008

2. Rahman MM, Teng J, Worrell BT, Noviello CM, Lee M, Karlin A, Stowell MHB, Hibbs RE (2020) Structure of the native muscle-type nicotinic receptor and inhibition by snake venom toxins. Neuron. doi: https://doi.org/10.1016/j.neuron.2020.03.012

3. Changeux J-P (2018) The nicotinic acetylcholine receptor: a typical ‘allosteric machine’. Philos Trans R Soc Lond 373 (1749):20170174. doi: 10.1098/rstb.2017.0174

4. Gharpure A, Noviello CM, Hibbs RE (2020) Progress in nicotinic receptor structural biology. Neuropharmacology 171:108086. doi: https://doi.org/10.1016/j.neuropharm.2020.108086

5. Griffon N, Büttner C, Nicke A, Kuhse J, Schmalzing G, Betz H (1999) Molecular determinants of glycine receptor subunit assembly. The EMBO journal 18 (17):4711–4721. doi: 10.1093/emboj/18.17.4711

6. Haeger S, Kuzmin D, Detro-Dassen S, Lang N, Kilb M, Tsetlin V, Betz H, Laube B, Schmalzing G (2010) An intramembrane aromatic network determines pentameric assembly of Cys-loop receptors. Nat Struct Mol Biol 17 (1):90–98. doi: 10.1038/nsmb.1721

7. Wong L-W, Tae H-S, Cromer BA (2014) Role of the ρ1 GABAC receptor n-terminus in assembly, trafficking and function. ACS Chem Neurosci 5 (12):1266–1277. doi: 10.1021/cn500220t

8. Pandhare A, Pirayesh E, Stuebler AG, Jansen M (2019) Triple arginines as molecular determinants for pentameric assembly of the intracellular domain of 5-HT3A receptors. J Gen Physiol 151 (9):1135–1145. doi: 10.1085/jgp.201912421

9. Yu X-M, Hall ZW (1991) Extracellular domains mediating ɛ subunit interactions of muscle acetylcholine receptor. Nature 352 (6330):64–67. doi: 10.1038/352064a0

10. Green WN, Claudio T (1993) Acetylcholine receptor assembly: Subunit folding and oligomerization occur sequentially. Cell 74 (1):57–69. doi: 10.1016/0092-8674(93)90294-Z

11. Blount P, Merlie JP (1990) Mutational analysis of muscle nicotinic acetylcholine receptor subunit assembly. J Cell Biol 111 (6):2613–2622. doi: 10.1083/jcb.111.6.2613

12. Merlie JP, Lindstrom J (1983) Assembly in vivo of mouse muscle acetylcholine receptor: Identification of an α1-subunit species that may be an assembly intermediate. Cell 34 (3):747–757. doi: 10.1016/0092-8674(83)90531-7

13. Saedi MS, Conroy WG, Lindstrom J (1991) Assembly of Torpedo acetylcholine receptors in Xenopus oocytes. J Cell Biol 112 (5):1007–1015. doi: 10.1083/jcb.112.5.1007

14. Gu Y, Forsayeth JR, Verrall S, Yu XM, Hall ZW (1991) Assembly of the mammalian muscle acetylcholine receptor in transfected COS cells. J Cell Biol 114 (4):799–807. doi: 10.1083/jcb.114.4.799

15. Kreienkamp H-J, Maeda RK, Sine SM, Taylor P (1995) Intersubunit contacts governing assembly of the mammalian nicotinic acetylcholine receptor. Neuron 14:635–644

16. Blount P, Smith MM, Merlie JP (1990) Assembly intermediates of the mouse muscle nicotinic acetylcholine receptor in stably transfected fibroblasts. J Cell Biol 111 (6):2601–2611. doi: 10.1083/jcb.111.6.2601

17. Charnet P, Labarca C, Lester HA (1992) Structure of the gamma-less nicotinic acetylcholine receptor: learning from omission. Mol Pharmacol 41 (4):708

18. Jackson MB, Imoto K Fau - Mishina M, Mishina M Fau - Konno T, Konno T Fau - Numa S, Numa S Fau - Sakmann B, Sakmann B (1990) Spontaneous and agonist-induced openings of an acetylcholine receptor channel composed of bovine muscle alpha-, beta- and delta-subunits. Euro J Physiol 417 (0031-6768 (Print)):129–135

19. Kullberg R, Owens JL, Camacho P, Mandel G, Brehm P (1990) Multiple conductance classes of mouse nicotinic acetylcholine receptors expressed in Xenopus oocytes. Proc Natl Acad Sci USA 87 (6):2067–2071. doi: 10.1073/pnas.87.6.2067

20. Park J-Y, Mott M, Williams T, Ikeda H, Wen H, Linhoff M, Ono F (2014) A single mutation in the acetylcholine receptor δ-subunit causes distinct effects in two types of neuromuscular synapses. J Neurosci 34 (31):10211–10218. doi: 10.1523/JNEUROSCI.0426-14.2014

21. Liu Y, Brehm P (1993) Expression of subunit-omitted mouse nicotinic acetylcholine receptors in Xenopus laevis oocytes. J Physiol 470 (1):349–363. doi: 10.1113/jphysiol.1993.sp019862

22. Nicke A, Rettinger J, Mutschler E, Schmalzing G (1999) Blue native page as a useful method for the analysis of the assembly of distinct combinations of nicotinic acetylcholine receptor subunits. J Recept SigTransduct 19 (1-4):493–507. doi: 10.3109/10799899909036667

23. Gu Y, Camacho P, Gardner P, Hall ZW (1991) Identification of two amino acid residues in the ϵ subunit that promote mammalian muscle acetylcholine receptor assembly in COS cells. Neuron 6 (6):879–887. doi: https://doi.org/10.1016/0896-6273(91)90228-R

24. Gharpure A, Teng J, Zhuang Y, Noviello CM, Walsh RM, Cabuco R, Howard RJ, Zaveri NT, Lindahl E, Hibbs RE (2019) Agonist selectivity and ion permeation in the α3β4 ganglionic nicotinic receptor. Neuron 104 (3):501-511.e506. doi: https://doi.org/10.1016/j.neuron.2019.07.030

25. Edgar RC (2004) MUSCLE: Multiple sequence alignment with high accuracy and high throughput. Nucleic Acids Research 32 (5):1792–1797

26. Webb B, Sali A (2014) Comparative protein structure modeling using modeller. Curr Prot Bioinf 5:5.61-65.66.32

27. Trott O, Olson AJ (2010) AutoDock Vina: improving the speed and accuracy of docking with a new scoring function, efficient optimization, and multithreading. J Comput Chem 31 (2):455–461. doi: 10.1002/jcc.21334

28. Gasteiger J, Marsili M (1978) New model for calculating atomic charges in molecules. Tetrahedron Lett 19 (34):3181–3184

29. Goodsell DS, Olson AJ (1990) Automated docking of substrates to proteins by simulated annealing. Proteins: Struct Function Bioinf 8 (3):195–202. doi: 10.1002/prot.340080302

30. Kandt C, Ash WL, Peter Tieleman D (2007) Setting up and running molecular dynamics simulations of membrane proteins. Methods 41 (4):475–488. doi: https://doi.org/10.1016/j.ymeth.2006.08.006

31. Jorgensen WL, Chandresekhar J, Madura JD, Impey RW, Klein ML (1983) Comparison of simple potential functions for simulating liquid water. J Chem Phys 79:926–935

32. Darden T, York D, Pedersen L (1993) Particle mesh Ewald - an N.log(N) method for Ewald sums in large systems. J Chem Phys 98 (12):10089–10092

33. Hess B (2008) P-lincs: A parallel linear constraint solver for molecular simulation. J Chem Theor Comput 4 (1):116–122. doi: 10.1021/ct700200b

34. Lindorff-Larsen K, Piana S, Palmo K, Maragakis P, Klepeis J, Dror R, Shaw D (2010) Improved side-chain torsion potentials for the Amber ff99SB protein force field. Proteins: Struc Func Genet 78:1950–1958

35. Jämbeck JPM, Lyubartsev AP (2012) Derivation and systematic validation of a refined all-atom force field for phosphatidylcholine lipids. J Phys Chem B 116 (10):3164–3179. doi: 10.1021/jp212503e

36. Berendsen HJC, Postma JPM, van Gunsteren WF, DiNola A, Haak JR (1984) Molecular dynamics with coupling to an external bath. J Chem Phys 81:3684–3690

37. Bussi G, Donadio D, Parrinello M (2007) Canonical sampling through velocity rescaling. J Chem Phys 126

38. Evans DJ, Holian BL (1985) The Nose–Hoover thermostat. J Chem Phys 83 (8):4069–4074. doi: 10.1063/1.449071

39. Abraham MJ, Murtola T, Schulz R, Páll S, Smith JC, Hess B, Lindahl E (2015) GROMACS: High performance molecular simulations through multi-level parallelism from laptops to supercomputers. SoftwareX 1–2:19–25. doi: http://dx.doi.org/10.1016/j.softx.2015.06.001

40. Monastyrskyy B, D’Andrea D, Fidelis K, Tramontano A, Kryshtafovych A (2016) New encouraging developments in contact prediction: Assessment of the CASP11 results. Proteins: Struct Func Bioinf 84 (S1):131–144. doi: 10.1002/prot.24943

41. Gowers RJ, Linke M, Barnoud J, Reddy TJE, Melo MN, Seyler SL, Dotson DL, Domanski J, Buchoux S, Kenney IM, Beckstein O MDAnalysis: A Python package for the rapid analysis of molecular dynamics simulations. In: Benthall S, Rostrup S (eds) Python in Science, Austin, Tx., 2016.

42. Heckman KL, Pease LR (2007) Gene splicing and mutagenesis by PCR-driven overlap extension. Nature Protocols 2 (4):924–932. doi: 10.1038/nprot.2007.132

43. Cetin H, Epstein M, Liu WW, Maxwell S, Rodriguez Cruz PM, Cossins J, Vincent A, Webster R, Biggin PC, Beeson D (2019) Muscle acetylcholine receptor conversion into chloride conductance at positive potentials by a single mutation. Proc Nat Acad Sci 116:21228–21235. doi: 10.1073/pnas.1908284116

44. Moroni M, Vijayan R, Carbone A, Zwart R, Biggin PC, Bermudez I (2008) Non-agonist-binding subunit interfaces confer distinct functional signatures to the alternate stoichiometries of the {alpha}4{beta}2 nicotinic receptor: An {alpha}4-{alpha}4 interface is required for Zn^2+^ potentiation. J Neurosci 28 (27):6884–6894

45. Nayak TK, Bruhova I, Chakraborty S, Gupta S, Zheng W, Auerbach A (2014) Functional differences between neurotransmitter binding sites of muscle acetylcholine receptors. Proc Natl Acad Sci USA

46. Hall ZW, Sanes JR (1993) Synaptic structure and development: The neuromuscular junction. Cell 72:99–121. doi: 10.1016/S0092-8674(05)80031-5

47. Brehm P, Henderson L (1988) Regulation of acetylcholine receptor channel function during development of skeletal muscle. Devel Biol 129 (1):1–11. doi: https://doi.org/10.1016/0012-1606(88)90156-X

48. Galzi J-L, Changeux J-P (1994) Neurotransmitter-gated ion channels as unconventional allosteric proteins. Curr Opin Struct Biol 4 (4):554–565. doi: https://doi.org/10.1016/S0959-440X(94)90218-6

49. Gross A, Ballivet M, Rungger D, Bertrand D (1991) Neuronal nicotinic acetylcholine receptors expressed in Xenopus oocytes: role of the α subunit in agonist sensitivity and desensitization. Pflügers Archiv 419 (5):545–551. doi: 10.1007/BF00370805

50. Couturier S, Erkman L, Valera S, Rungger D, Bertrand S, Boulter J, Ballivet M, Bertrand D (1990) Alpha 5, alpha 3, and non-alpha 3. Three clustered avian genes encoding neuronal nicotinic acetylcholine receptor-related subunits. J Biol Chem 265 (29):17560–17567

51. Chavez-Noriega LE, Crona JH, Washburn MS, Urrutia A, Elliott KJ, Johnson EC (1997) Pharmacological characterization of recombinant human neuronal nicotinic acetylcholine receptors hα2β2, hα2β4, hα3β2, hα3β4, hα4β2, hα4β4 and hα7 expressed in *Xenopus* oocytes. J Pharm Exp Thera 280 (1):346

52. Unwin N (2005) Refined Structure of the Nicotinic Acetylcholine Receptor at 4Å Resolution. J Mol Biol 346:967–989

53. Rahman MM, Teng J, Worrell BT, Noviello CM, Lee M, Karlin A, Stowell MHB, Hibbs RE (2020) Structure of the native muscle-type nicotinic receptor and inhibition by snake venom toxins. Neuron:in press. doi: 10.1016/j.neuron.2020.03.012

54. Mazzaferro S, Benallegue N, Carbone A, Gasparri F, Vijayan R, Biggin PC, Moroni M, Bermudez I (2011) An additional ACh binding site at the α4/α4 interface of the (α4β2)2α4 nicotinic acetylcholine receptor confers stoichiometry-specific properties. J Biol Chem 286:31043–31054

55. Prevost MS, Bouchenaki H, Barilone N, Gielen M, Corringer P-J (2020) Concatemers to re-investigate the role of α5 in α4β2 nicotinic receptors. Cell Mol Life Sci:in press. doi: 10.1007/s00018-020-03558-z

56. Jansen M, Bali M, Akabas MH (2008) Modular design of cys-loop ligand-gated ion channels: Functional 5-HT3 and GABA ρ1 receptors lacking the large cytoplasmic M3M4 loop. J Gen Physiol 131 (2):137–146. doi: 10.1085/jgp.200709896

57. Kukhtina V, Kottwitz D, Strauss H, Heise B, Chebotareva N, Tsetlin V, Hucho F (2006) Intracellular domain of nicotinic acetylcholine receptor: the importance of being unfolded. J Neurochemistry 97 (s1):63–67. doi: 10.1111/j.1471-4159.2005.03468.x

58. Kuhse J, Laube B, Magalei D, Betz H (1993) Assembly of the inhibitory glycine receptor: Identification of amino acid sequence motifs governing subunit stoichiometry. Neuron 11 (6):1049–1056. doi: https://doi.org/10.1016/0896-6273(93)90218-G

59. Jha A, Cadugan DJ, Purohit P, Auerbach A (2007) Acetylcholine receptor gating at extracellular transmembrane domain interface: the cys-loop and M2-M3 linker. J Gen Physiol 130 (6):547–558. doi: 10.1085/jgp.200709856

60. Lee WY, Sine SM (2005) Principal pathway coupling agonist binding to channel gating in nicotinic receptors. Nature 438 (7065):243–247. doi: 10.1038/nature04156

61. Grosman C, Salamone FN, Sine SM, Auerbach A (2000) The extracellular linker of muscle acetylcholine receptor channels is a gating control element. J Gen Physiol 116 (3):327–340. doi: 10.1085/jgp.116.3.327

62. Jensen ML, Schousboe A, Ahring PK (2005) Charge selectivity of the Cys-loop family of ligand-gated ion channels. J Neurochemistry 92 (2):217–225. doi: 10.1111/j.1471-4159.2004.02883.x

63. Oliveira ASF, Edsall CJ, Woods CJ, Bates P, Nunez GV, Wonnacott S, Bermudez I, Ciccotti G, Gallagher T, Sessions RB, Mulholland AJ (2019) A general mechanism for signal propagation in the nicotinic acetylcholine receptor family. J Am Chem Soc 141 (51):19953–19958. doi: 10.1021/jacs.9b09055

64. Akk G, Sine S, Auerbach A (1996) Binding sites contribute unequally to the gating of mouse nicotinic alpha D200N acetylcholine receptors. J Physiol 496 (Pt 1) (Pt 1):185–196. doi: 10.1113/jphysiol.1996.sp021676

