## Supplementary Information for "Structural correlates of human muscle nicotinic acetylcholine receptor subunit assembly mediated by δ(+) interface residues"

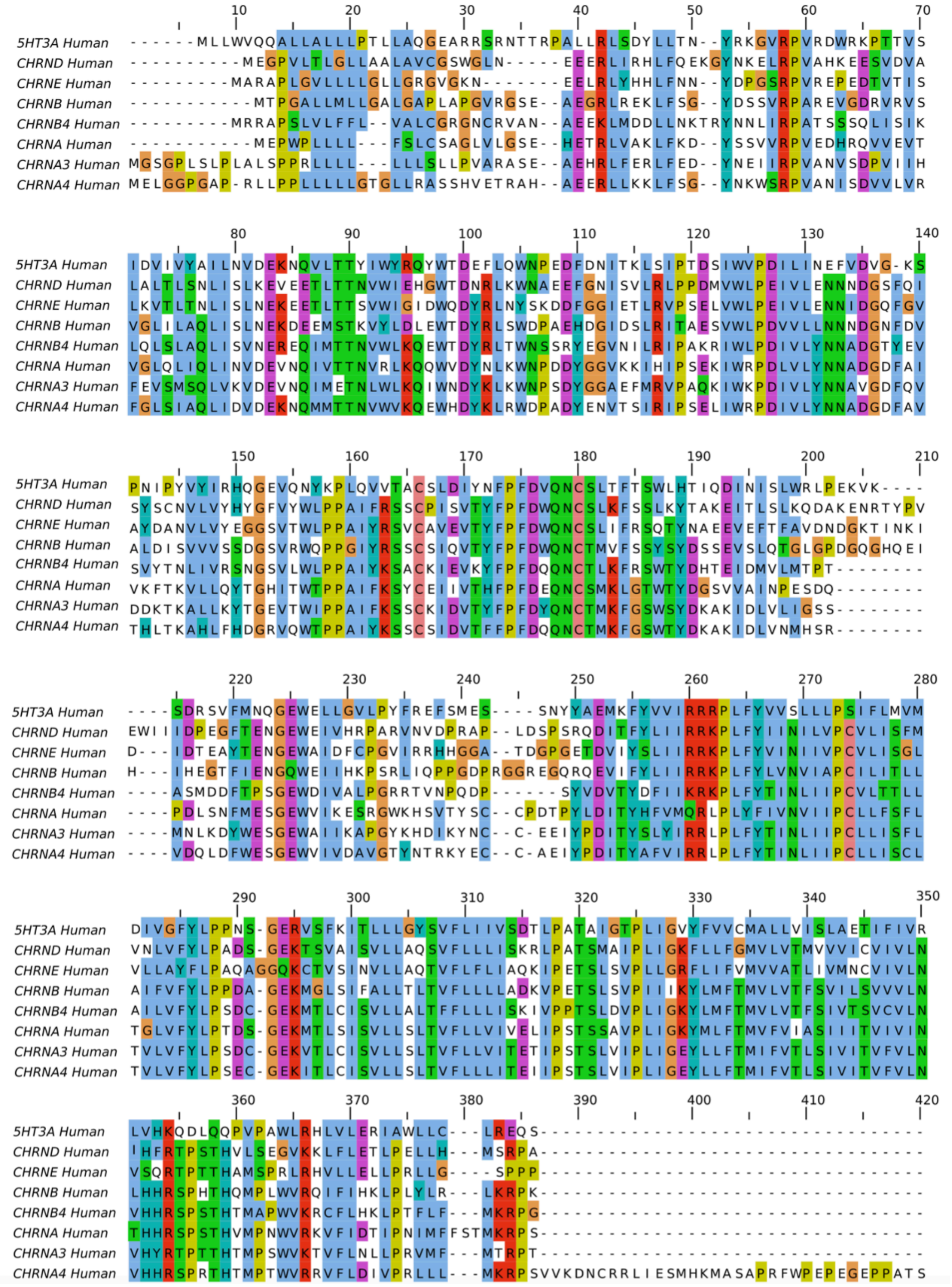
5HT3A Human
CHRND Human
CHRNE Human
CHRNB Human
CHRNB4 Human
CHRNA Human
CHRNA3 Human
CHRNA4 Human

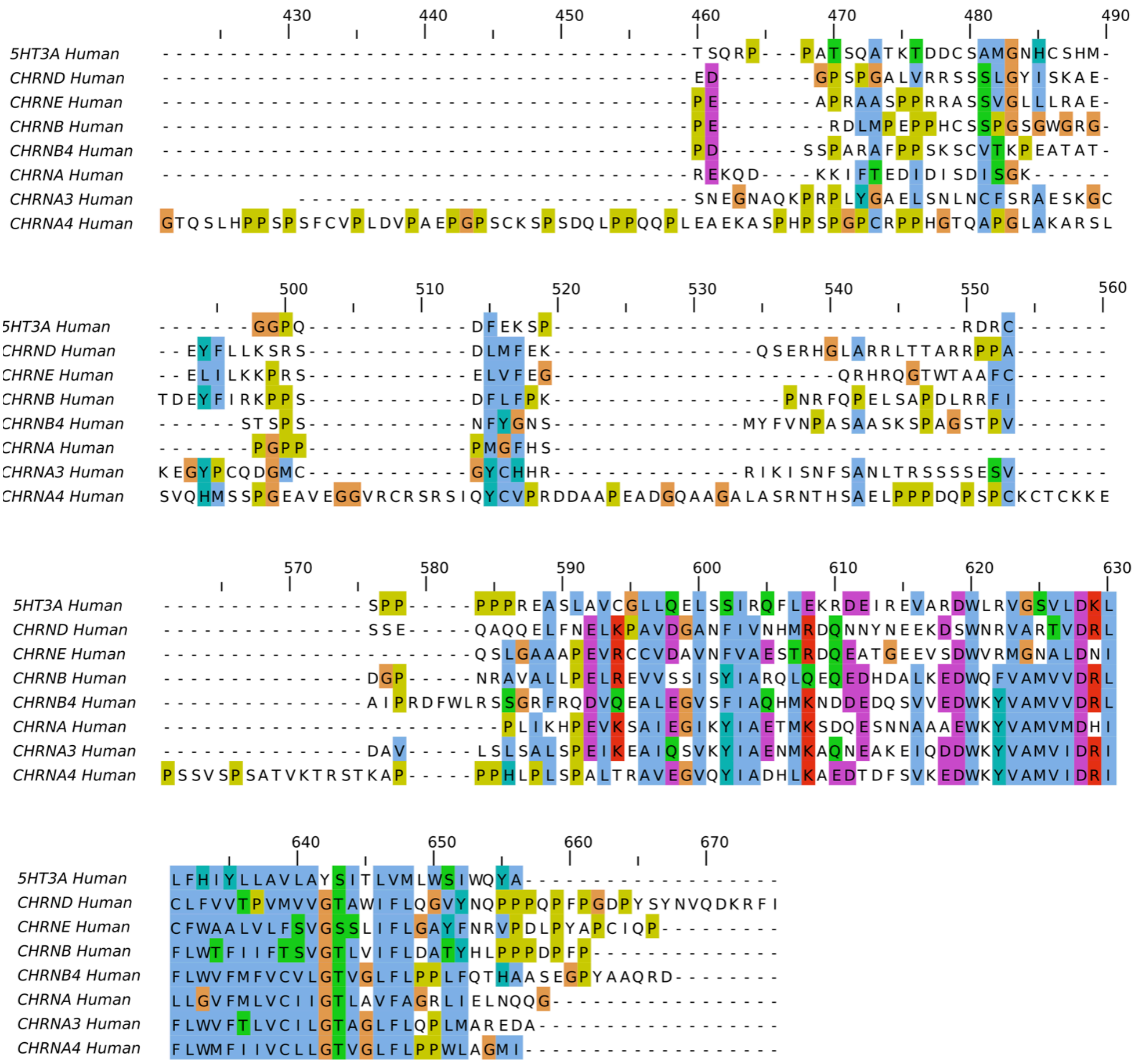


**SI Figure. 1.** Multiple sequence alignment used to generate the initial model.

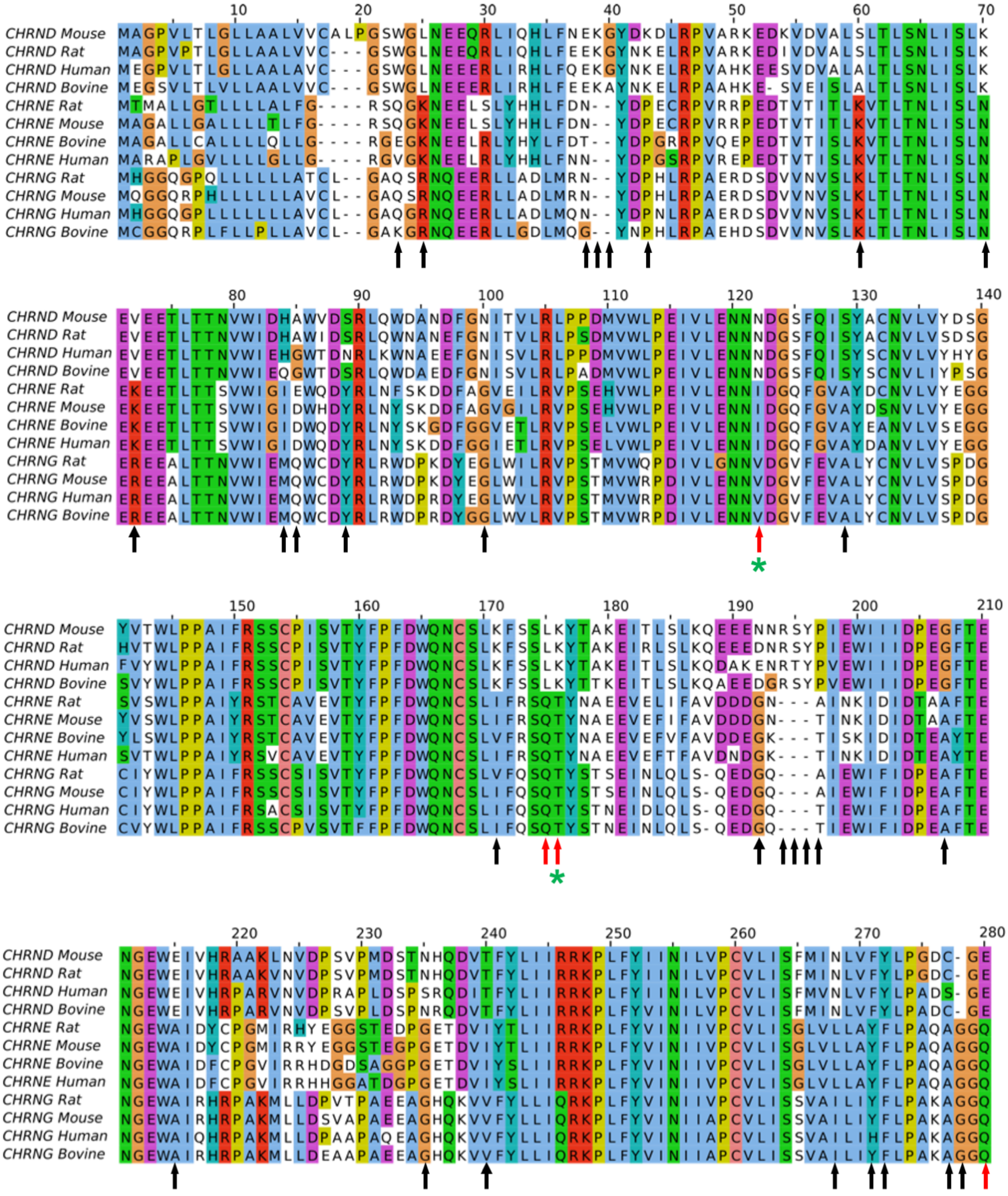
CHRND Mouse
CHRND Rat
CHRND Human
CHRND Bovine
CHRNE Rat
CHRNE Mouse
CHRNE Bovine
CHRNE Human
CHRNG Rat
CHRNG Mouse
CHRNG Human
CHRNG Bovine
10
20
30
40
50
60
70
80
90
100
110
120
130
140
150
160
170
180
190
200
210
220
230
240
250
260
270
280

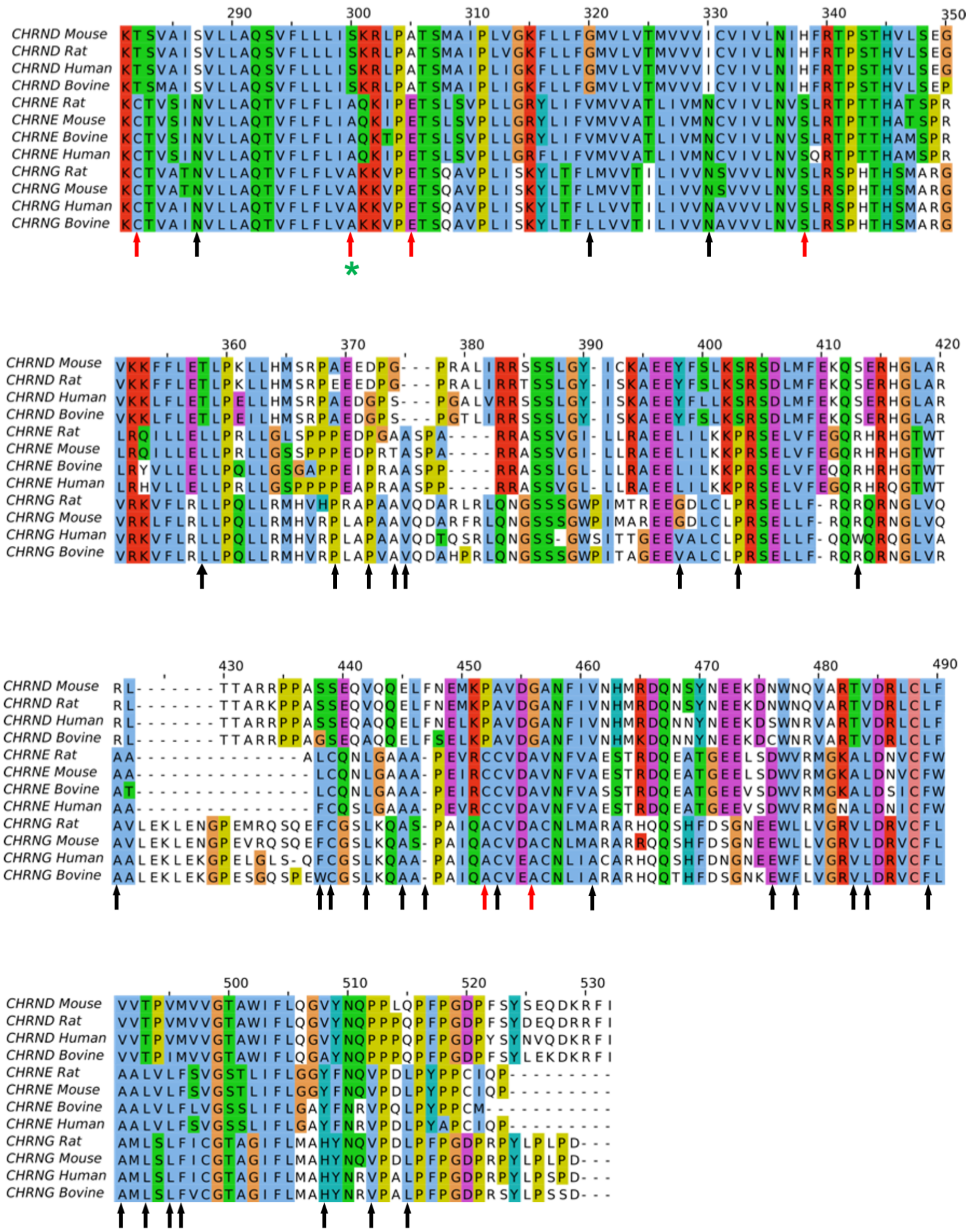


**SI Figure. 2.** Multiple sequence alignment of mammalian δ, ε and ɣ subunits. Black arrows highlight residues where ɣ and ε are conserved, but where δ is not. Red arrows indicate residues that match this evolutionary criteria and also where the corresponding δ subunit Cβ atoms falls within 8 Å of a neighbouring β subunit interface

Cβ atom. Green asterixis highlights the position of the residues forming contacts that are analyzed in Fig. 4 of the main manuscript.

**SI Table 1. Primers used for overlap extension.**

| | |
|---|---|
| Forward epsilon 5' δ overlap | ATCATCAACATCCTGGTGCCCTGTGTGCTCATCTCG |
| Reverse epsilon 5' δ overlap | CGAGATGAGCACACAGGGCACCAGGATGTTGATGAT |
| BamHI restriction with Kozak sequence. δ forward | TATGGATCCGCCACCATGGCTGAAATGGAGGGG |
| BamHI restriction with Kozak sequence. δ forward | AGCTCTAGACTAAGGCTGGATACACGGCGC |

**SI Table 2**. δ-residues that were identified from the contact matrix and MSA integration with % of frames of which their respective β-carbons were within 8 Å of β-face β-carbons. Percentages over 100 indicate contacts with multiple residues on the β-face.

| Residues | % Contact of aggregate simulation time |
|---|---|
| N119 | 287 |
| L172 | 291 |
| K173 | 260 |
| E277 | 120 |
| T279 | 406 |
| S297 | 170 |
| A302 | 365 |
| H335 | 73 |
| P439 | 179 |
| G442 | 365 |

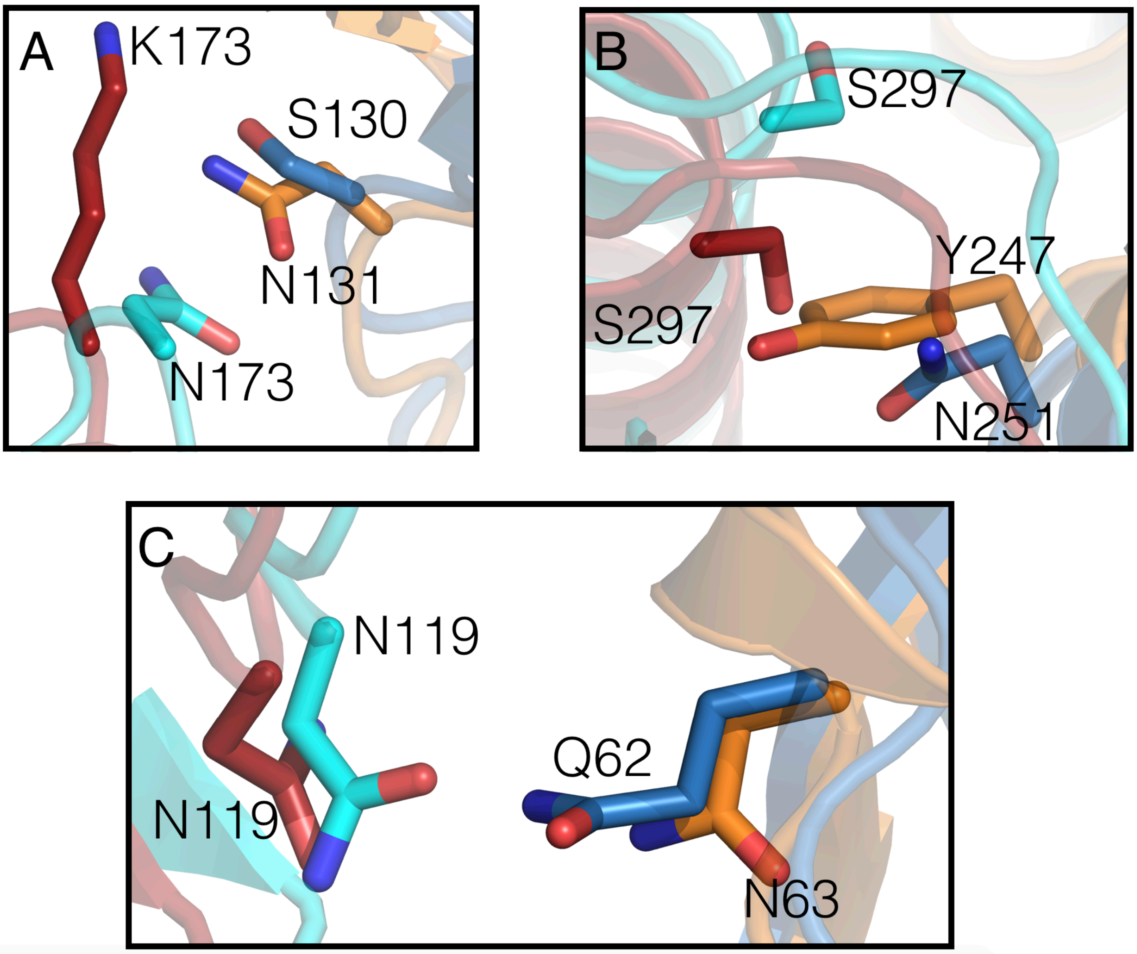


**SI Figure 3.** Superimposition of adult human muscle nAChR comparative model δ & β subunits coloured in red and blue on *Torpedo* 6UWZ δ and β subunits coloured in cyan and orange respectively. The RMSD was 2.43 Å (Pymol Align tool). Panels A, B and C represent identified assembly regions K173, S297 and N119. Only residue N119 is conserved across *Torpedo* and human receptors.

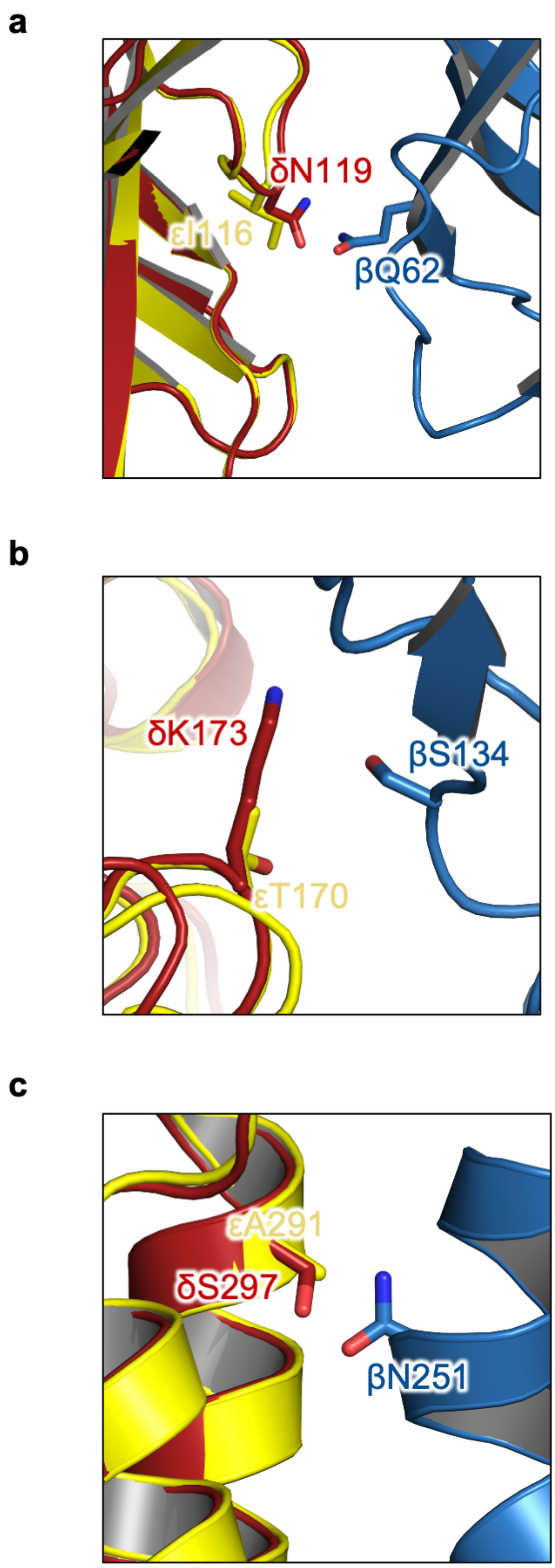


**SI Figure 4** Superimposition of ε (yellow) on δ (red) subunit with neighbouring β (blue) subunit. The homologous residue to δN119 on the ε subunit is isoleucine 116 (A). The homologous residue to δK173 on the ε subunit is threonine 170 (B) and the homologous residue to δS297 on the ε subunit is alanine 291

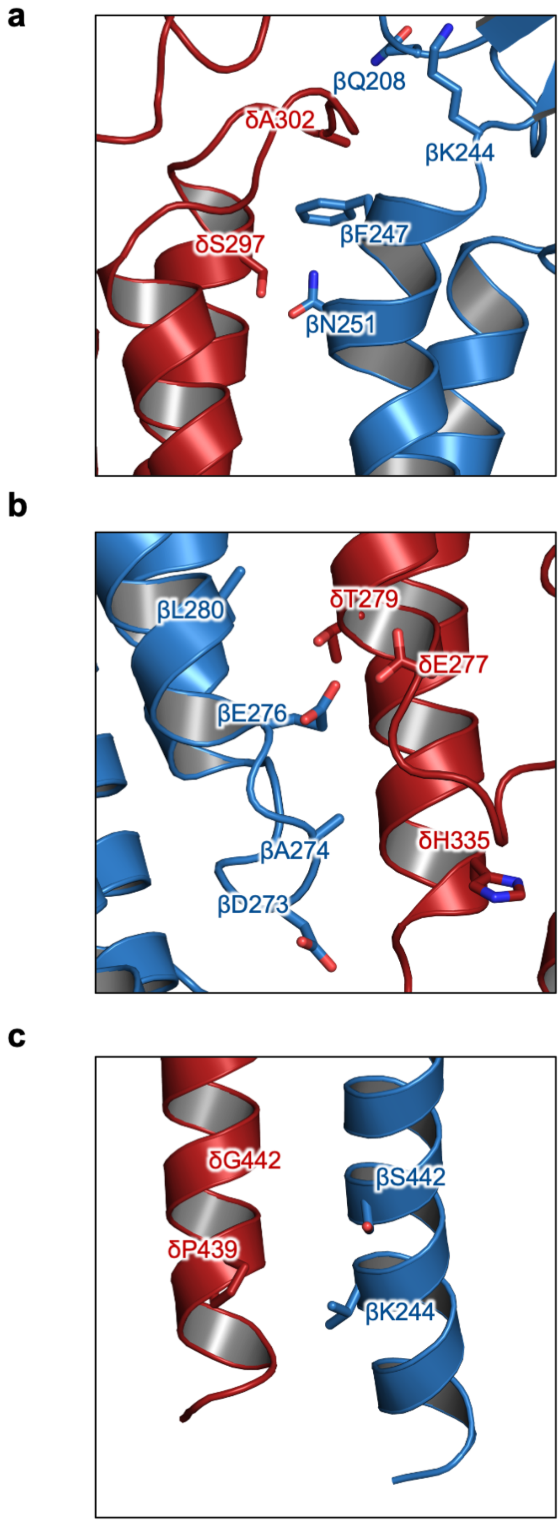


**SI Figure 5** Diagramatic summary of the locations of TMD contacts made as summarized in SI Table 2. (A) δA302 is located at the top of the TMD and sits between βF247 and the alkyl sidechain of βK244. (B) δE277 likely faces the interior of the pore (viewed flipped 180°). (C) δP439 and δG442 are located towards the intracellular end.

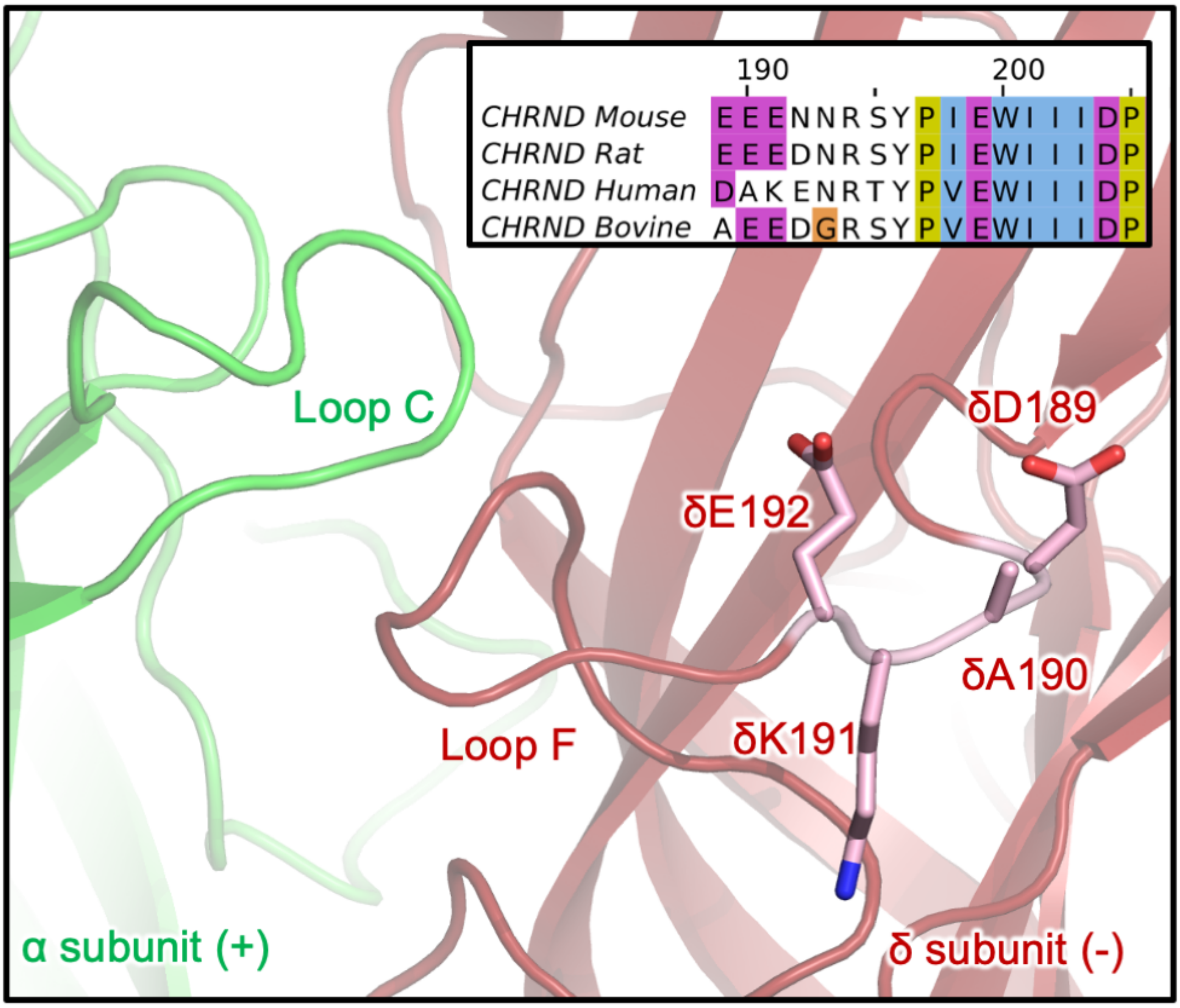


**SI Figure 6.** Snapshot of αδ interface extracellular domain Subunits are shown in cartoon representation with α and δ subunits in green and red respectively. Non-conserved δ subunit residues on Loop F (as shown in the sequence alignment insert) are shown in pink stick representation.
